# Representational geometry explains puzzling error distributions in behavioral tasks

**DOI:** 10.1101/2023.01.03.522667

**Authors:** Xue-Xin Wei, Michael Woodford

## Abstract

Measuring and interpreting errors in behavioral tasks is critical for understanding cognition. Conventional wisdom assumes that encoding/decoding errors for continuous variables in behavioral tasks should naturally have Gaussian distributions, so that deviations from normality in the empirical data indicate the presence of more complex sources of noise. This line of reasoning has been central for prior research on working memory. Here we re-assess this assumption, and find that even in ideal observer models with Gaussian encoding noise, the error distribution is generally non-Gaussian, contrary to the commonly held belief. Critically, we find that the shape of the error distribution is determined by the geometrical structure of the encoding manifold via a simple rule. In the case of a high-dimensional geometry, the error distributions naturally exhibit flat tails. Using this novel insight, we apply our theory to visual short-term memory tasks, and find that it can account for a large array of experimental data with only two free parameters. Our results call attention to the geometry of the representation as a critically important, yet underappreciated factor in determining the character of errors in human behavior.

## 1 Introduction

Human cognition is prone to errors [1, 2, 3, 4, 5, 6]. Such errors range from profound perceptual illusions to more subtle errors that only become apparent in carefully designed behavioral experiments. Despite that such errors may be undesirable in various circumstances, they could help elucidate the basic question of how the brain performs computations [7, 8, 9, 10, 11, 12, 13, 14, 15, 16, 17, 18].

One dogmatic assumption in the studies of human behavior has been that the error distribution of a continuous variable should a *priori* be Gaussian distributed. This assumption is critical, because it is reasoned that any deviations of the error distributions from Gaussianility should be indicative of the existence of additional mechanisms. This line of reasoning has profound implications in the field of visual short-term memory (VSTM) [3, 19, 20, 21], which widely employs continuous estimation paradigms to investigate brain computations. Many studies reported that error distributions for multiple stimulus attributes (e.g., orientation, color) deviates substantially from Gaussianality [17, 22, 23, 24, 25, 26], often exhibiting flat long tails. Such non-Gaussianality has been interpreted as evidence in favor of a fixed capacity limit [3, 27, 28, 20, 29, 17, 22, 23, 24], variability in encoding precision [25, 26, 30], dynamical allocation of memory resources [31, 32], variability of samples from the neural spiking process [33, 34, 35], and more recently nonlinear psychological scaling [36], and is still under heavy debated to date.

Despite the popularity of this belief of Gaussianality, its validity has not been explicitly examined. This strong belief is possibly due to the intuition that adding noises from many independent sources would effectively result in Gaussian noise. However, this argument has a major gap. That is, Gaussian measurement noise may not necessarily lead to Gaussian error distribution, as the structure of the signal may also matter. To address this question, we study the error distributions in Bayesian observer models, which provide optimal estimate given an encoding model and have been used to explain various aspects of cognition [7, 8, 9, 10, 13, 11, 37, 38]. Surprisingly, we find that ideal observer models generally predict non-Gaussian error distributions even with Gaussian measurement noise, contradicting the conventional wisdom. Critically, we elucidate that how the geometry of the representation acts as the primary factor that determines the shape of the error distribution. This theory provides accurate account of a wide array of experimental data with two, or even fewer parameters. Our results demonstrate the importance of studying the geometry of the neural representation and its relation to behavior.

## 2 Results

Consider a visual perception or memory task, in which the subject needs to report the stimulus (e.g., a particular color on the color wheel) they just saw. To model a task of this kind, we assume the information of a particular stimulus attribute *θ* is encoded by a neural representation *R* via an encoding model *p*(*R*|*θ*). Subsequently, the reported stimulus attribute by the observer corresponds to a readout of this representation by inverting the encoding model to obtain a posterior distribution and the Bayesian estimate according to Bayes rule. This general Bayesian framework has been applied to model many behavioral tasks. One key component of this approach, and an active area of research, is the structure of the neural representation *p*(*R*|*θ*). Previous studies have specified it via neural tuning curves [39, 40, 41, 42], signal detection theory framework [43, 14], efficient coding [15, 44, 45, 16, 46], or other characteristics of the neural codes [13].

### 2.1 Bayesian observer model based on representational geometry

We assume that the representation of stimulus variable *θ* forms a response manifold embedded [47, 48] in a space with sufficiently large dimensionality. The representation of stimuli *θ*, denoted as *m*(*θ*), is assumed to be corrupted by homeostatic Gaussian noise with magnitude *σ* along each dimension.

We next define the response manifold by directly specifying its representational geometry. The representational distance (RD) *d*(*θ_i_*, *θ_j_*) between each pair of stimuli *θ_i_* and *θ_j_* is calculated as the Euclidean distance between their representations *m*(*θ_i_*), *m*(*θ_j_*). RD determines the stimulus discriminability [48], thus is well motivated behaviorally. We further assume the representation is symmetric: first, the RD only depends on the stimulus disparity, i.e. *d*(*θ_i_*, *θ_j_*)^2^ = *g*(*θ_i_* – *θ_j_*), where *g* is a dissimilarity function that satisfies *g*(0) = 0; second, the manifold has a centroid, and its distance to *m*(*θ*) is 1 for every stimulus. For a circular variable such as orientation or color on a color wheel, the stimulus disparity is computed by computing arc distance on the circle. Note that it is possible to specify the model using tuning curves. In this case, we have

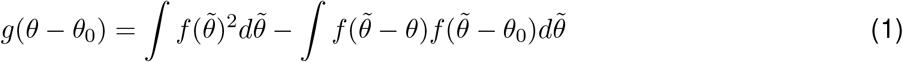

where *f*(*θ*–*θ*_0_) is the tuning curve for a neuron with preferred stimulus *θ*_0_. However, we prefer to describe the model directly in terms of its geometry (i.e., the dissimilarity function *g*) as this is more generic; that is, the same representation geometry may correspond to many different tuning configurations (see SI Fig. S1) that all lead to the same error distribution.

We model the behavioral response of each trial using the maximum a posteriori probability (MAP) estimate 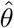. Assuming a uniform prior over *θ*, it is equivalent to computing the stimulus that maximizes the log-likelihood function, which essentially quantifies the Euclidean distance between the random measurement and the mean representation of each stimulus. Thus, finding the Bayesian estimate amounts to identify whose representation is closest to the measurement. If the model is specified using tuning curves as above, we find that the log-likelihood function (treated as a random function) can be expressed as

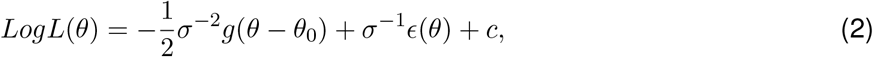

where the true stimulus is denoted as *θ*_0_, and *c* is constant (see Methods for derivations). The first component in the log-likelihood is a fixed component determined by the RD. It may be interpreted as quantifying the overall strength of a“matching signal” between the memory representation and the elicited measurement, which is determined by the dissimilarity. The second component is a random component *ϵ*(*θ*) due to the fluctuation of the measurement around *m*(*θ*_0_). Because noise will lead to correlated change of the distance to different stimuli, therefore the second component is correlated, and critically, the correlation structure can also be compactly described by the representational geometry. More precisely, here *ϵ*(*θ*) is a homogeneous Gaussian random field with zero mean, and 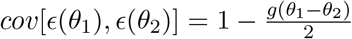. Thus both the “matching signal” and the correlation structure of the “matching signal” are determined by the representational dissimilarity function *g*. Note that the tuning curve *f*(.) doesn’t actually matter, other than for the geometry that it implies.

### 2.2 Geometry determines the shape of the error distribution

To model circular variable such as orientation and color on a color wheel, we take *f* as a von Mises function with a concentration parameter *κ*, and its peak magnitude ensures 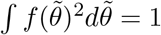. The *κ* parameter determines the dissimilarity function thus the geometry (see SI Fig. S2). With this parameterization, the entire model only has two free parameters, the geometry parameter *κ* and the noise parameter *σ*. Both parameters are highly interpretable. While here we will focus on this simple parameterization, other more complicated parameterizations are possible. For instance, one could specify f as a weighted sum of von Mises functions (see SI fig S3), resulting in a more flexible family of dissimilarity functions with more parameters.

Parameter *κ* in our model determines the dimensionality of the representation. A higher *κ* value implies a representation of higher dimensionality. A convenient way to evaluate the embedding dimensionality is through a principal component analysis (PCA). As *κ* is made larger, the first two principal dimensions account for less of the total variance; conversely, more dimensions are needed to accounted for 95% of the variance of the mean response manifold (see SI Fig. S4). The effect of *κ* on the geometrical structure of the manifold can be visualized intuitively via classic multidimensional scaling (MDS) analysis. When the *κ* is close to 0, the MDS embedding closely resembles a circle. As *κ* becomes larger, the manifold becomes increasingly curved, as seen from a 3-dimensional MDS visualization.

Fig. 1 shows the error distribution corresponding to three different encoding geometries. Somewhat surprisingly, the error distributions appear to have a variety of shapes, ranging from Gaussian-like to those with flat tails. Interestingly, plotting the logarithm of error density in the similarity scale revealed an approximately linear relationship for all three cases (which becomes exact in the case of a linear manifold).

**Figure 1:**
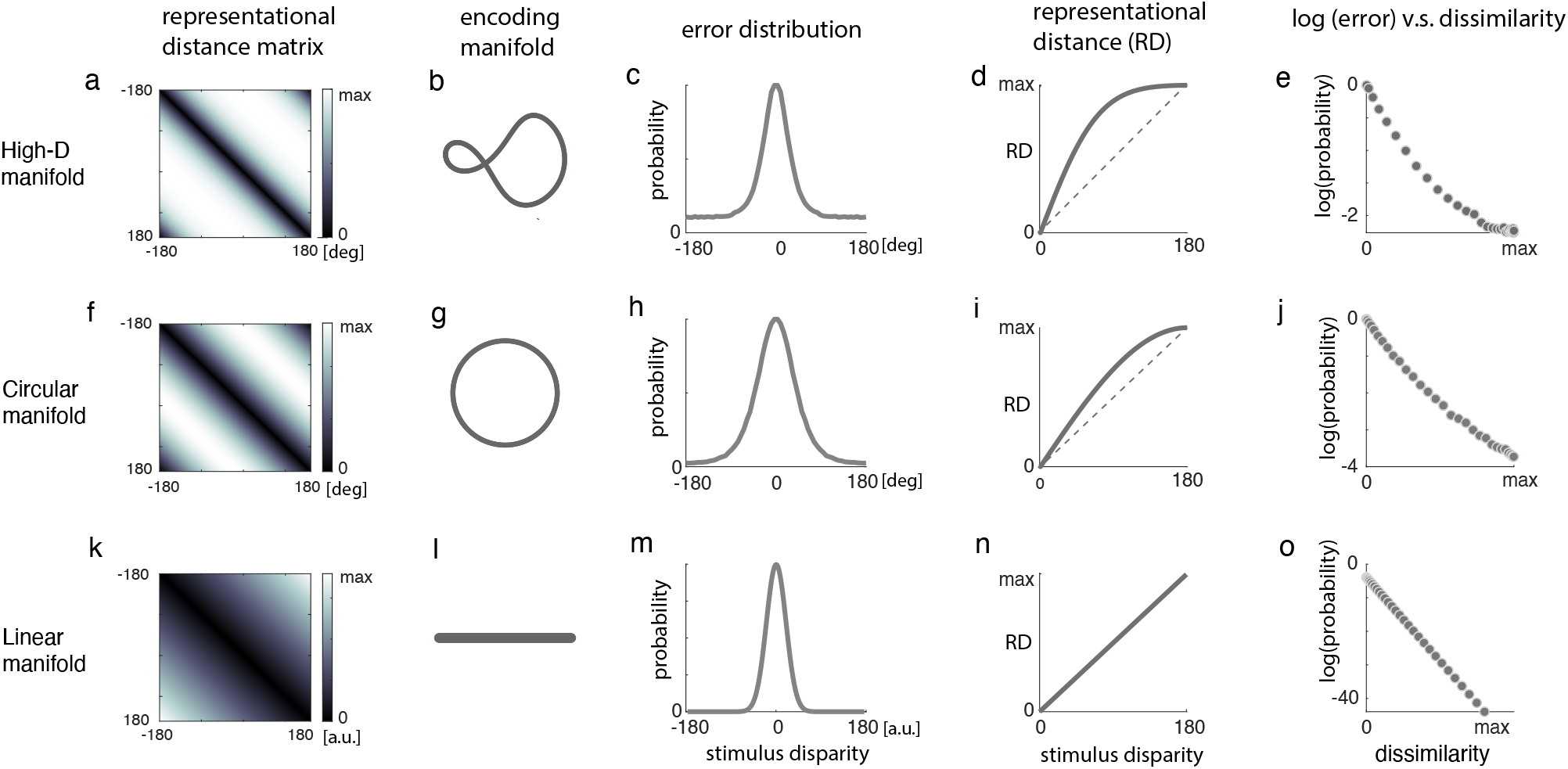
Different manifold geometry leads to different shapes of error distributions. (a-e) When encoding manifold is high dimensional, the error distribution exhibits a heavy and even close to flat tail. (a) Representation distance matrix that specifies the geometry. (b) A visualization of the encoding manifold using classic multi-dimensional scaling in 3-D. (c) Example error distribution when simulating the proposed Bayesian observer model. The heavy tail is apparent. (d) The representational distance (RD) as a function of the stimulus disparity. RD shows an early saturation. (e) When plotting the logarithm of the error density function in the dissimilarity scale, the two show an approximately linear relationship. (f-j) Similar to (a-e), but for a circular manifold. In this case, the tail of the error distribution is not as heavy as that for the high-D manifold. Again, when plotting the logarithm of the error density function in the dissimilarity scale, the two show an approximately linear relationship. (k-o) Similar to (a-e), but for Linear manifold. In this case, the quantities shown can be computed analytically. With a linear encoding manifold,the error distribution will be Gaussian. This is consistent with the traditional wisdom of Gaussian error distribution[17]. However, in general, error distribution will not be Gaussian, and its shape depends on the geometry of the encoding.

To understand the characteristics of the error distributions more systematically, we varied the parameters *σ*, *κ* in our model. This investigation further consolidates the observation made in Fig. 1e,i,o. That is, despite of the diverse shapes of the error distributions in the stimulus space, to the first order the error density decays exponentially when considering the similarity (or equivalently, dissimilarity) scale defined by *g*(.). This holds across different encoding geometries and noise levels. Due to the quadratic relationship between the representational distance (RD) and the dissimilarity function *g*(.), this also means that the error should decay as a Gaussian function of the representational distance (Fig. 2gj). This observation provides a novel theoretical insight regarding how geometry of the representation determines the outcome of the Bayesian inference. It is striking that the shape of the error distribution becomes straightforward to interpret when the geometry is considered.

**Figure 2:**
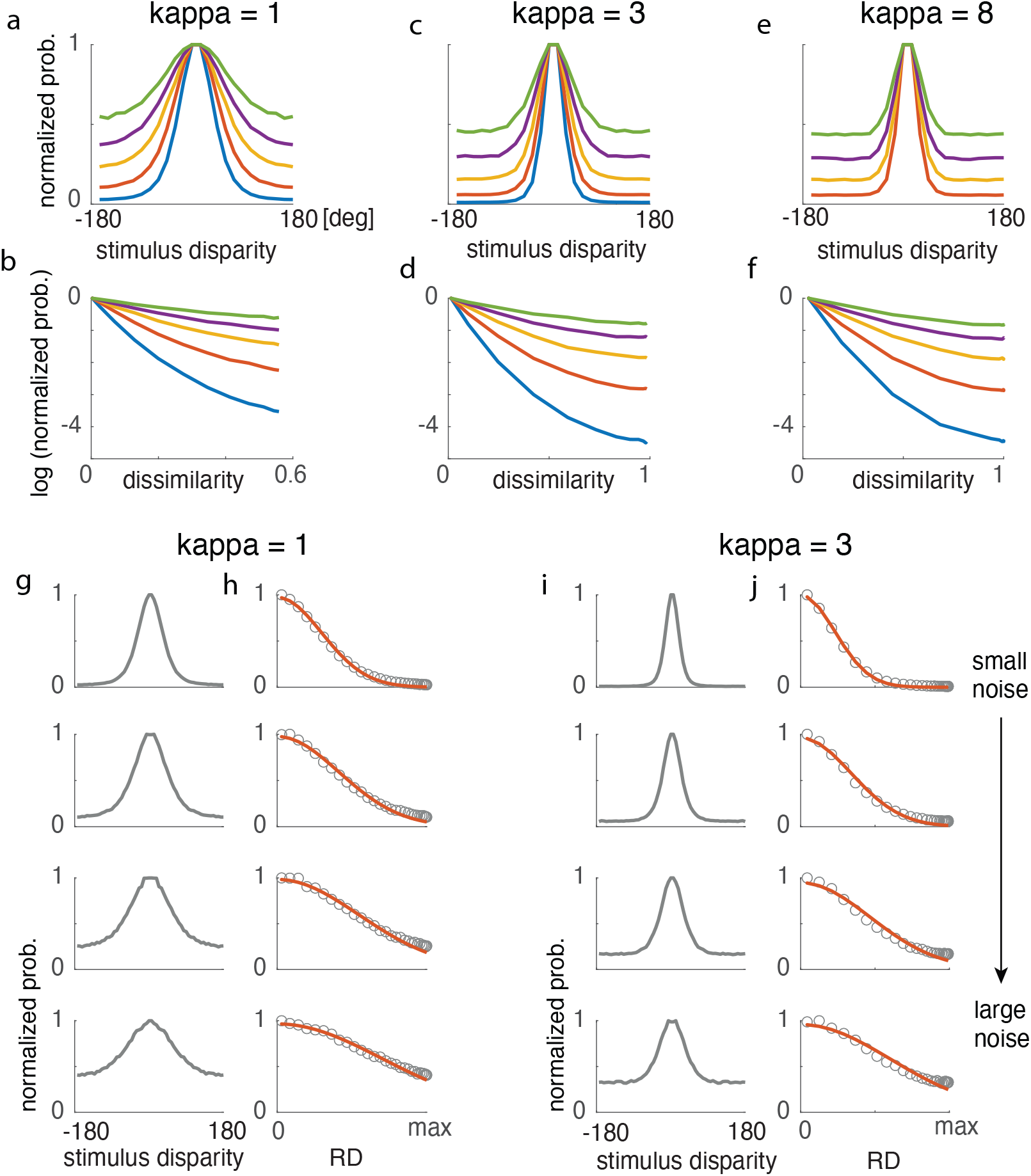
The model predicts that the error density function should decay approximately as a Gaussian function in the representational distance scale. (a,b) The logarithm of the error density roughly follow a linear relationship with the representational dissimilarity function. Notice the slight deviation from linearity for large dissimilarity. (c,d,e,f) Similar to (a,b), but for two other *κ* values. (g,h) The error density (shown in g), when considered in the representational distance scale(RD, i.e., the square root of dissimilarity function *g*()), can be well approximated by a Gaussian function (shown in h). This holds for different noise levels and geometries. Red curves plots the Gaussian fit to the warped error density functions in the RD scale. The Gaussian fits, while not perfect, capture the warped error density function.

For a given stimulus *θ*_0_, the probability that the reported memory item being *θ*_0_ + *δθ* can be approximated as

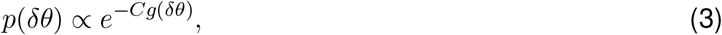

where *C* is a constant that depends on the noise *σ*. It is worth emphasizing that this relationship is not exact, in particular when the error is large. This inexactness could be seen through the subtle deviation from linearity in Fig. 2b,d,f, which plot the logarithm of the error density (i.e., *logp*(*δθ*)) as a function of the similarity function *g*(*δθ*) under different noise levels. Despite of the slight inaccuracy, this relationship remains useful in understanding the shape of the error distribution, as it gives a simple description of the relation between the representational geometry and the error distribution.

Eq. 3 implies that the density functions of the error under different amount of noise should be linked by a power function. Indeed, we found that that we could indeed use the error density under medium noise condition together with a power function to accurately predict error distributions under much larger or smaller noise levels (see Fig. S5).

Importantly, the two parameters in our model have qualitatively distinct consequences on the error distribution. First, *σ* affects how fast the error probability decays via the constant *C*(*σ*), without much change of its shape (e.g., heaviness of the tail). Here *C*(*σ*) is an increasing function of sigma. The relationship between 1/*σ* and C is roughly linear and varies depending on the geometry (see Fig. S6). In contrast, *κ* controls the shape of the error distribution through a change of the similarity function (Fig. 2a-f). Thus, the parameters in the model are highly interpretable.

Previously, the impact of the encoding geometry on the error distributions has essentially been ignored. It was often assumed a *priori* that the error should decay as a Gaussian function in the stimulus space.^1^ Such an assumption is reasonable if the neural manifold is flat, in which case the Guassian noise would lead to Gaussian error distributions. However, our results demonstrated that, generally speaking, the error distribution should be non-Gaussian, and the geometry of the encoding heavily influences the shape of the error distribution. Curved manifold naturally leads to deviation from Gaussianality. With larger *κ*, the representation becomes increasingly curved and high dimensional (Fig. S4), thus representational distance saturates at smaller stimulus disparity(Fig. S2), leading to a long and flat tail of the error distribution.

When does Gaussian measurement noise lead to Gaussian error distribution then? Our theory identifies two such scenarios: first, the encoding manifold straight (i.e.,without extrinsic curvature; see Fig 1k-o); second, the noise is sufficiently small to make the estimation problem local, in which case the relevant part portion of manifold is approximately straight.

### 2.3 The model accounts for error patterns in memory tasks

Over the past several decades, studies of VSTM have accumulated a large amount of data concerning the pattern of errors in various types of tasks, in particular estimation tasks (e.g., [51, 17, 32, 52, 50, 25, 26, 33, 24, 23, 53, 36, 45]), that thus provide a good testbed for our theory..

#### Color data

We start by examining whether our model could explain the data from several color VSTM experiments [17, 32, 25, 36], which employed a continuous report paradigm. In each trial the observer needs to identify the color of one particular previously seen items on a color wheel. A key experimental variable that is manipulated is set size, which often ranges from 1 to 8. To model the effect of the set size or length of delay, we assume they only affect the noise *σ*, not the geometry of the encoding (i.e., parameter *κ*). Although this is likely a simplified assumption as it does not directly model the potential interference between different items, below we show that it already accurately accounts for the pattern of a large set of data.

We fit the color data in [36] jointly, with each condition (e.g., different set sizes or lengths of delay) has its own noise parameter, while share the geometry parameter. This results in 17 parameters in total for 16 datasets. These color data across the different conditions in [36] could be well fitted by our model (Fig. 3a). This includes the data from conditions with large set sizes, for which the tails of the error distributions appear to be flat. Previously, such flat tails have motivated the development of slotbased model [17], which proposes that the flat tails in the error distributions reflect guesses due to the limited capacity of working memory systems. Here we found that the flat tail emerges naturally when the encoding manifold is high-dimensional, due to that the similarity function saturates at large stimulus disparity, thus offers a conceptually very different explanations.

**Figure 3:**
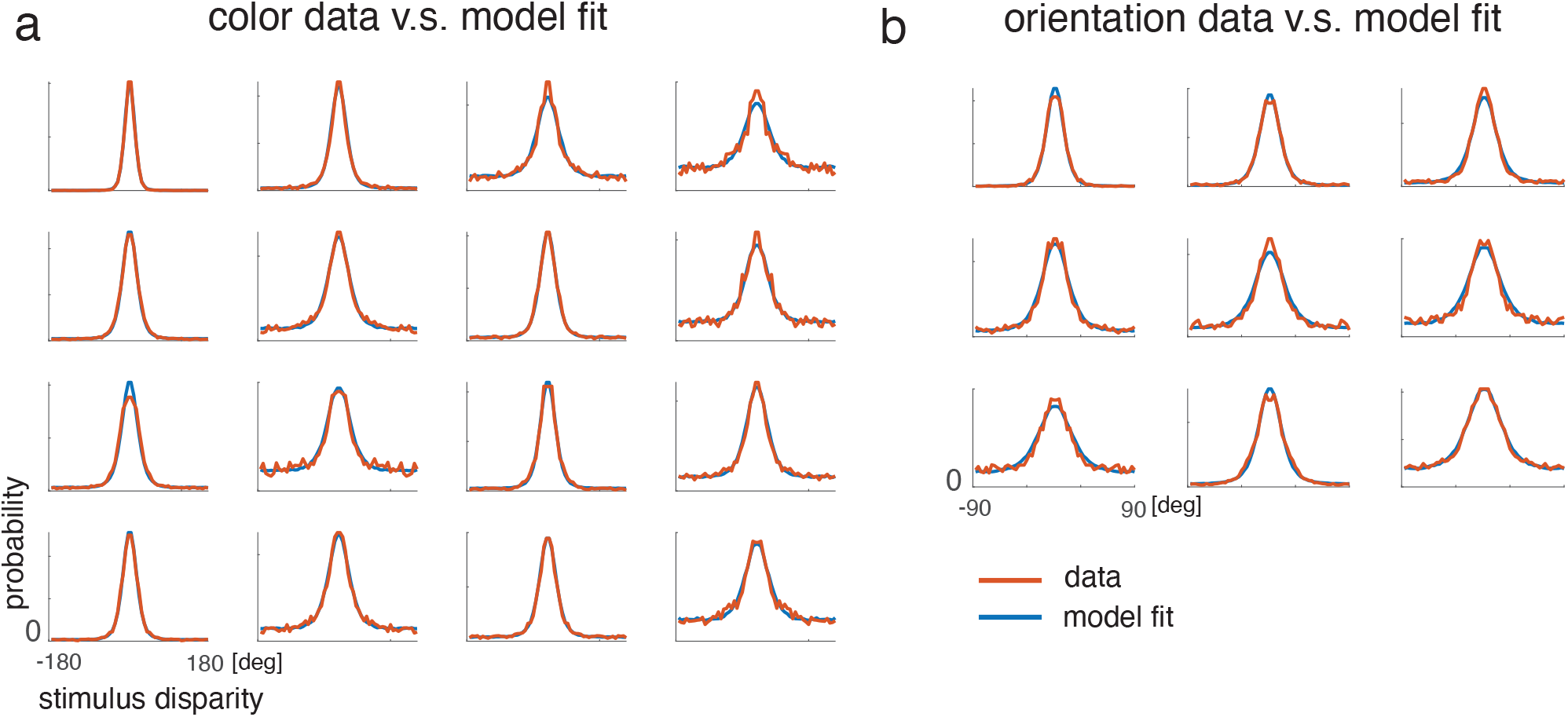
The model with two free parameters can well fit the VSTM data from color and orientation experiments. a) Color data from [36], and model fit. Here, we fit the 16 conditions jointly. The *kappa* parameter is shared, while the noise parameter *σ* is private to each condition. Thus, overall there are 17 free parameters for these 16 conditions. b) Orientation data from [25] and [50], and model fit. Again, these 9 conditions are jointly fitted with 10 free parameters in total.

Using the same value of *κ* parameter inferred above, we further performed a single parameter fit to other color data reported in several other color VSTM experiments (24 conditions) [17, 32, 25]. Again, our model is sufficient for fitting these empirically measured error distributions (Fig. S7).

The theory predicts that warping the error density function using the representation distance (RD) scale should result in approximately a Gaussian function. Fig. 4a shows that indeed the warped the density function is approximately Gaussian for most of the conditions. A related prediction is that the logarithm of error distribution should decay linearly in the dissimilarity scale (Fig. 4b). To test this, we plotted the logarithm of the measured error distributions against the dissimilarity function inferred. The relationship between the two is well described by a linear model.

**Figure 4:**
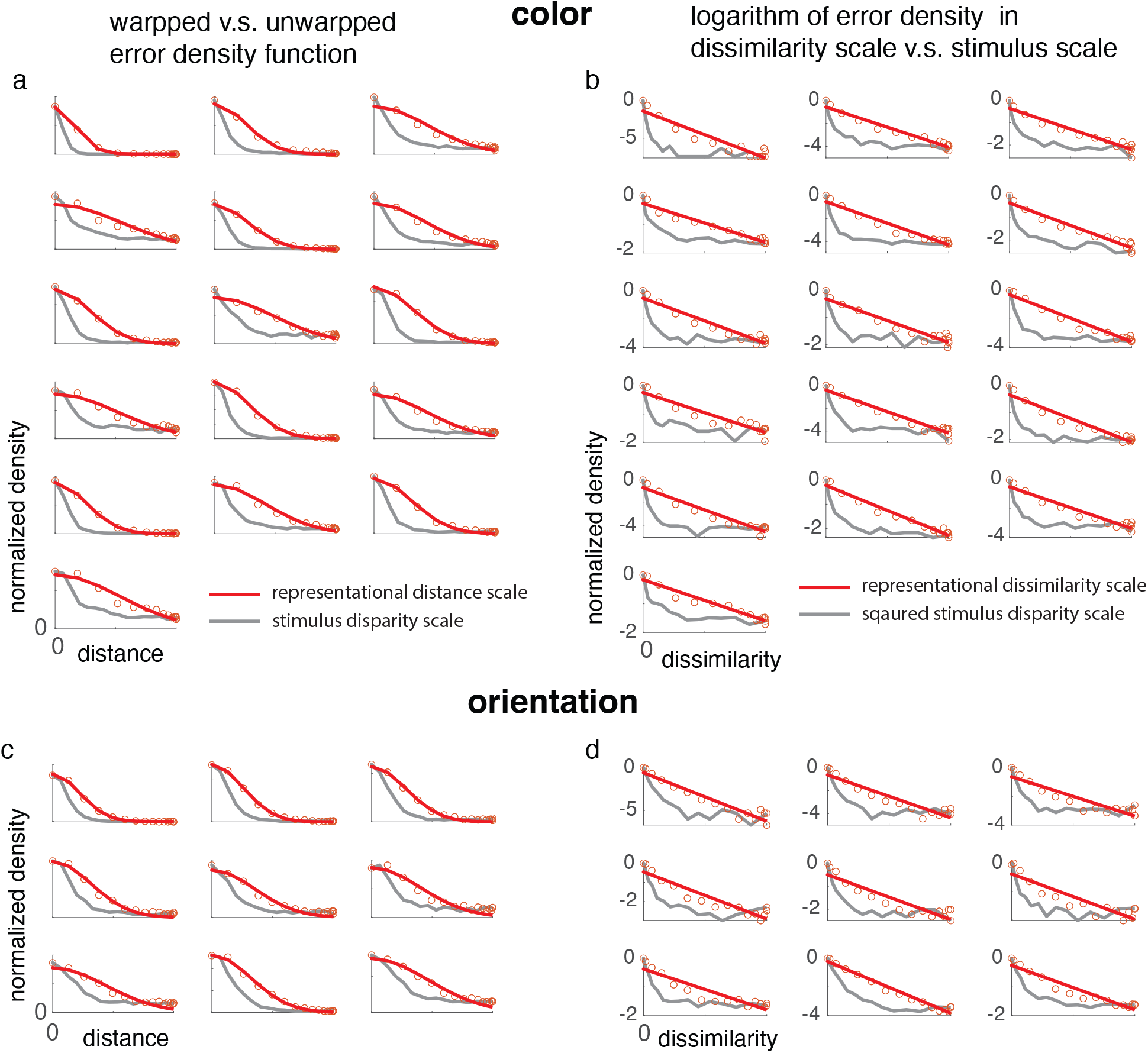
Testing the model prediction that the error density function should decay approximately as a Gaussian function in the representational distance (RD) scale.(a) Comparison of the warped and the unwarped error density function for the color VSTM data. Each panel represents one experimental dataset. The grey line represents the error density function plotted in the original stimulus scale. The red dots re-plot the error density function in the (warped) representational distance scale. After warping, the error density function is close to a Gaussian function. The red line is a Gaussian fit to the red dots. (b) Grey curves: the logarithm of the error density (normalized to its maximum) plotted against the squared stimulus distance along the circle. Red dots: re-plotting the logarithm of the error density function in the (warped) dissimilarity scale. After warping, the logarithm of error density function is approximately a linear function of the dissimilarity. Red lines: Linear fits to the red dots. A perfect fit would imply that the error density function decays exactly as a Gaussian function in the representational distance (RD) scale. Data from [36] (c,d) Similar to (a,b), but for orientation data. Data are from [25, 50].

#### Orientation data

To test whether the same model account for other stimulus attributes, we also fit our model to datasets from orientation([50]) VSTM tasks. These datasets (9 conditions included in total) were collected using similar procedure as the color data above. The results (Fig. 3b) show that the model provides strong fits to those data, suggesting the applicability of our theory is not specific to color. Using warping analysis via the dissimilarity scale inferred, we again found strong evidence supporting our theory (Fig. 4c,d).

#### Comparison between color and orientation

Interestingly, the inferred geometries for orientation and color are different. The best-fitting *κ* parameters for the color and orientation data are 3.4, and 1.6 respectively. This implies that the error distributions for color end to have flatter tails than those for color. This difference naturally arises in our framework as the orientation and color may exhibit different encoding geometry. It may be difficult to explain this difference in some previous models, including slot models [17] and variable precision models [25, 49], which ignore the specific encoding properties of the each stimulus modality.

#### n-AFC data

According to our model, the only factor that could change the overall performance for a given stimulus attribute is the noise level *σ*, thus the performance in a 2-AFC task should be sufficient to predict error distribution when more options are available (e.g., 360-AFC or continuous estimation). [36] collected data using color stimuli from the same set of observers to perform AFC tasks for color with varying number of options. Leveraging these data, we inferred the noise level for each observer based the 2-AFC task. Using the geometry parameter inferred from the continuous report task above, we generate zero-free-parameter predictions of the error distribution for 60-AFC and 360-AFC tasks. Results in Fig. 5 show that the predicted error density well matches the measurement from the 360-AFC task and 60-AFC task. The ability to accurately predict the n-AFC task from the 2-AFC data without any additional parameter provides a strong test of our model. Note that a recent model described in [36] also well predicted the n-AFC data based on the 2-AFC data, however, additional assumptions on the noise correlation structure were needed in that model.

**Figure 5:**
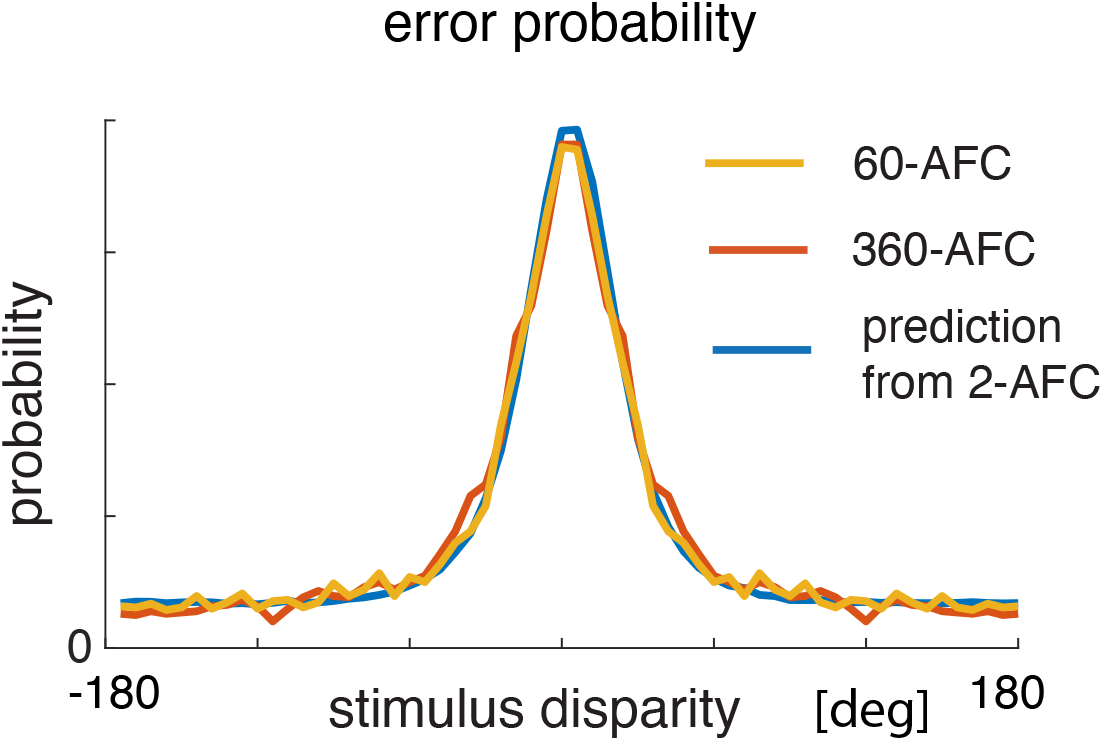
Using our model, one can well predict the data 60-AFC (orange) task and 360-AFC (red curve) task from the 2-AFC data. Blue: prediction for the continuous estimation task (360-AFC) from the 2-AFC data using our model. The predicted error distribution appears to be slightly peakier than the measured, yet the overall shape of the measured error distribution is well captured by the prediction. The experimental data were collected in [36].

Together, these results demonstrate that our model with only two free parameters is sufficient to account for the data reported in various VSTM tasks across different stimulus domains, providing strong support of our model.

## 3 Discussion

We have studied the characteristics of error distributions and demonstrated the importance of representational geometry. We found that, depending the encoding geometry, error distributions in ideal observer models exhibit markedly different shapes. Our new approach helps distill the essential factors that determined the characteristics of error distributions. Focusing on the application to data from VSTMt tasks, we found that this principled framework with two free parameters could readily account for the reported error distributions across many different experiments, including those exhibit an almost flat tail.

It is useful to connect and contrast our model with other work. We first consider population coding model for VSTM tasks, in which Poisson spiking noise was proposed to be the key reason for the heavy tail in error distributions [33, 54, 35, 34]. One version of the population coding model (assuming equally spaced von Mises tuning curves and Gaussian noise) can be seen as a discrete realization of our model. A more detailed discussion is given in the SI. Different from claims of the population coding models, we propose that the geometry is a more fundamental primitive that determines the shape of error distributions. This is a new insight that was not realized previously. The same geometry and thus error distribution could correspond to very different tuning curve configurations. Our theory also provides a much simpler and unified understanding of previous results on neural population coding-based approach to study VSTM. Notably, [35] proposed that the width of the tuning curve provides an upper bound of encoding precision. While related, it does not establish a explicit connection between the geometry of the representation and the shape of the error distribution.

A recently proposed psychological scaling model (i.e.,TCC model [36]) was shown to well account for patterns of errors in various VSTM tasks. In that model, on every trial a matching signal is generated for each possible option with its strength determined by a psychological scaling function corrupted by Gaussian additive noise. The observer’s choice is assumed to reflect the option with the largest matching signal. Recent work has attempted to bridge the TCC model and population coding model [55], though without establishing a clear relation. We found the structure of TCC model formulation is formally similar ours, except that the structure of the noise correlation of the matching signals. Compared to our model, TCC models have more parameters to be specified. In [36], most of these parameters were specified by additional behavioral tasks, e.g., from the triad or quad task. Our model only has two parameters, thus is more compact both in theory and in practice. One could constrain the representational geometry by fitting the model to data collected in one condition, and then perform a single parameter fit to other conditions. Practically, this is straightforward and does not require another set of measurement.

Many previous studies (*e.g*.,[17, 25, 49]) have assumed Gaussian errors as a baseline assumption. In the slot-mixture model, the heavier tail in error distributions (relatively to Gaussian) have been previously attributed to random guesses[17], while others have attributed it due to the variability in encoding [52, 25]. This mixture model has been adopted in many prior studies with various clinical applications for extracting the precision of the encoding and guess rate. Our results suggest that it may be misleading to interpret the width of the error distribution observed in large set size (often extracted by fitting a mixture model) as a measure of precision. Instead, we propose that the width parameter extracted there for large set sizes may instead reflect the geometry of the encoding. To be clear, our model does not rule out the possibility of guesses. However, our results suggest that when fitting the slot-mixture model or other types of mixture models [17, 52, 25], it would be important to modify the assumption of Gaussianality for the “non-guess” trials. Our results thus invite a revisit of these findings with a revised baseline assumption to take into account the representational geometry. Future high resolution data with large sample sizes within individual observers are likely needed for further separating the different models. Current data are limited in the sense that the stimulus-specific variability [23, 45] of an individual observer and the variability across observers are ignored. This makes it difficult to faithfully compare different models quantitatively given these mis-specified assumptions.

A number of future paths could be taken to generalize the current results. First, we have assumed, for convenience, that the neural representation has a perfect symmetry and it further assumes the encoding precision at each location is uniform, which is an assumption commonly made in previous studies (but see [23, 45]). Second, we assume a uniform prior. It should be possible to relax these assumptions by incorporating various kinds of asymmetry and by combining Bayesian inference with the efficient coding framework [56, 15, 44, 16, 45]. Third, we assumed a Euclidean geometry, generalization to other types of geometry [57, 58] would be an interesting future direction. Finally, it is also possible to configure the geometry by using more sophisticated models, *e.g*., by using a deep neural network model. We hypothesize that the geometry will still play a key role in determining the estimation biases in a much broader class of models.

It is worth emphasizing that our theory is general, and not specific to the VSTM tasks. For instance, it may be used to model perception or long-term memory. In fact, perceptual estimation, working memory, long-term memory may be unified within the same framework. If these different processes shared a common neural encoding manifold, the shape of the error distribution associated with each should also be shared [59]. Our theory thus may provide general insights in linking the geometrical structure of neural code and the behavior under various cognitive processes.

## Acknowledgement

We thank the authors of all the previous studies, from which the data re-analyzed here were collected, for making their data publicly available and making the current study possible. We thank Michael Hahn, Nikolaus Kriegeskorte and Paul Bays for helpful discussions.

## Methods

### Model formulation

We parameterize the neural manifold using the representational geometry. We assume that the representation is symmetric such that the representation distance between two arbitrary stimuli is only determined by their stimulus disparity, which is the shortest distance along the circle for a circular variable. We parameterize the representational dissimilarity function using the following form: 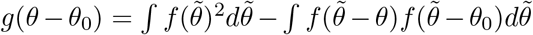. For modeling circular variable such as orientation, we take *f*(*θ*) = *Ae*^*κcos*(*θ*)^, which is a von Mises function with a concentration parameter *κ*. The normalization factor *A* ensures 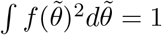. From this parameterization, it is straightforward to see that *g*(0) = 0.

We assume the representation of the stimulus is corrupted by an additive, homeostatic Gaussian noise. We denote the standard deviation of the noise along each dimension as *σ*. Thus, overall the model only has two free parameters, the geometry parameter *κ* and the noise parameter *σ*. A uniform prior is assumed throughout. The subject’s behavioral report on each trial is modeled as the stimulus value that maximizes the posterior distribution (i.e., MAP estimate), and equivalently, the logarithm of the posterior. The logarithm of the posterior essentially reduces to the log-likelihood function, which can be expressed as a random function that has the following form: 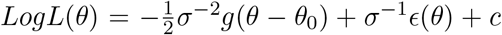. where the noise term *ϵ*(*θ*) is modeled by a homogeneous Gaussian random field, which is completely characterized by its first and second moments, *E*[*ϵ*(*θ*)] = 0, and 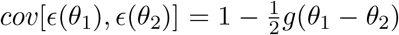. The final term *c* is a constant over *θ*.

### Deriving the log-likelihood function

Below, we derive the log-likelihood function based on our model. Denote the presentation for stimulus *θ* as *m*(*θ*), and is corrupted by homeostatic Gaussian noise *δ* with magnitude *σ* along each dimension. Note that both *m*(*θ*) and *δ* are random vectors. Denoting the centroid of the manifold as *m_c_*. Based on our assumptions, the representational distance between each stimulus and the centroid equals 1, *i.e*., *d*(*θ*, *m_c_*)^2^ = 1.

The log-likelihood function can be computed as

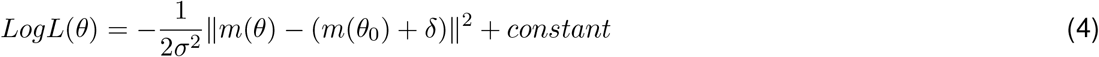

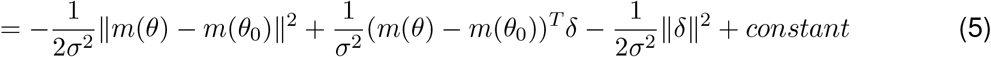

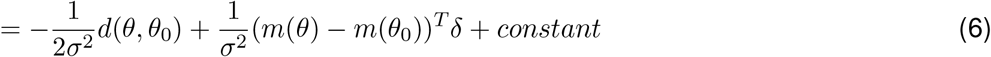

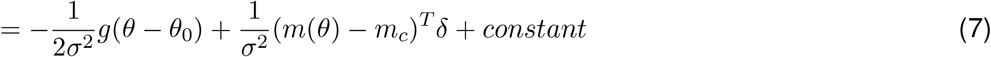

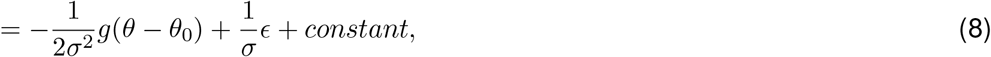

where, in the last step, we define 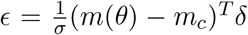. It is easy to verify that *E*(*ϵ*) = 0. Furthermore, the covariance structure can be derived as

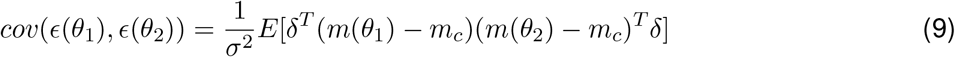

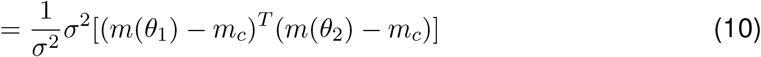

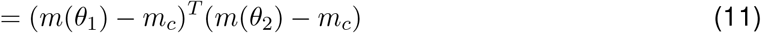

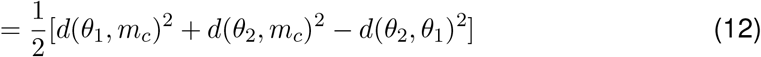

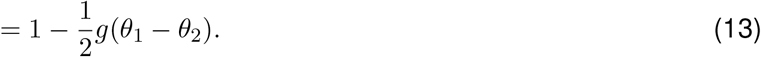

### Simulation

We discretized the stimulus points into 360 equally spaced grid pints. Given the geometrical configuration (specified by *κ*) and a noise level (specified by *σ*), the logarithm of the posterior is fully captured by a 360-dimensional multivariate Gaussian distributions. Obtaining the error distribution numerically is straightforward. We randomly generated multivariate Gaussian samples according to Eq. 2. More concretely, we sampled from a scaled version of the Log-likelihood function, −*g*(*θ* – *θ*_0_) + 2*σϵ*. The mean of the Gaussian is determined by −*g*(*θ_i_* – *θ*_0_) for each *θ_i_*, and the element of the covariance matrix between the *i^th^* and *j^th^* element is 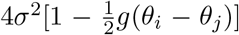. For each multivariate sample, we found the stimulus corresponding to the largest element. We varied the two parameters *κ* and *σ* in the model, and generated predicted error distributions for each combination of the two parameters. For each condition, we simulated a large number of trials, and convert the estimation errors into a predicted error density.

### Analyzing the error distributions predicted by the Bayesian model

We find that the predicted error distribution approximately decays exponentially as a function of the representational distance (RD). This is revealed by plotting the logarithm of the error density as a function of RD. We find that a powerlaw function could link the error density at different noise levels to each other. From the error density predicted based on an intermediate noise level (i.e., reference density in Fig. S5), we used the mean square error as the objective function to find the best matching exponent for each other noise level. We found that this single parameter fitting procedure is sufficient to predict error error densities under a large range of noise levels. We performed this analysis for multiple geometries (*κ* = 1, 3, 8), spanning the range relevant for the experiments based on our analysis of the previous experimental data.

### Modeling fitting to the data

Model fitting to the data was done using standard maximum likelihood estimation. We first numerically generated error distributions using numerical simulations for different combinations of *κ* and *σ*. The geometry parameter *κ* is taken to be have values equally spaced between 1 to 5, with a spacing of 0.2. The noise parameter *σ* is taken to have values between 2^-4^ and 1, evenly spaced on the logarithmic scale with a spacing of 0.1. This choice of parameter values ensures that we cover the relevant range based on our analysis of the experimental data. For each parameter set, we simulated 100000 samples to obtain the simulated error distribution. Given the measured error distribution, we evaluated the log-likelihood for each combination of parameters (*κ*,*σ*), and select the parameters that maximize the log-likelihood.

### Datasets

All datasets used in this paper were previously published and publicly available.

#### Color

We used (i) the datasets collected and reported in [36], and (ii) four datasets from the benchmark datasets collected from multiple labs, downloaded from https://www.cns.nyu.edu/malab/resources.html (labeled as Dataset #2,3,7,8 in the benchmark dataset). These four datasets were originally collected in references [17, 32, 25]. We refer the readers to the original publications for the details of the experiments. Briefly, in these experiments, a set of different colors drawn from a color wheel were presented in the presentation phase. After a delay period, the observers were asked to report the stimulus attribute for one or more items which they just saw on the color wheel. Set size was a key variable manipulated in these experiments. Different experiments have different collections of set sizes. For example, in [25], there are set sizes varying from 1 to 8. To calculate the error distributions for each set size, we pooled the data across different observers in the same experiments. For [17], we included set sizes 1,2,3,6. For [32], we included set sizes 1,2,4,6. For the experiments in [25] (Datasets #7,8 in the benchmark dataset), we included set size 1,2,3,4,5,6,7,8.

#### Orientation

The orientation data was also from the aforementioned benchmark datasets (labeled as Dataset 9, 10). The original data were collected in ref [25, 50]. In these experiments, orientation was used as the stimulus attributed, otherwise, the experiments were conducted similar to what was described above. For data from [25], we included set sizes 2,3,4,5,6,7,8. For data from [50], we included set sizes 3 and 6.

### Predicting the error distribution in n-AFC tasks from 2-AFC measurements

This analysis is based on the data collected in [36]. In their experiment, the same set of observers performed both a 2-AFC task and a 360-AFC task (a or 60-AFC task) with color (see [36] for details). We consider the 60-AFC task and 360-AFC task to be essentially continuous estimation tasks. We simulated the 2-AFC task under a range of noise parameter *σ* (specifically, we used 1000 noise levels that were equally spaced between 0.1 and 20) and evaluated the 2-AFC d-prime. This enables us to construct a lookup table that links the 2-AFC d-prime and the noise parameter *σ* in our model. In this dataset, the average accuracy across all observes is about 0.82, with substantial variability across observers. For each subject (n=52), we calculated the 2-AFC d-prime, then mapped onto the noise parameter. For the observers with a perfect performance, we set the 2-AFC d-prime to be 3. Using this set of inferred noise parameters and the fitted geometry parameter from datasets collected in [34] for color *κ* = 3.4, we generated a zero free-parameter prediction of the error distribution for the 360-AFC task by simulating our model using number of trials comparable to the experiments in [36].

## Footnotes

1 This is the basis of the mixture model [17] and variable precision model [25, 49].

