## Supplementary information for "Representational geometry explains puzzling error distributions in behavioral tasks"

### Supplementary notes: relationship of our model to other models

Our model could be re-formulated using other forms and connected to other models. The notes below further clarify the similarity and the difference of these models.

**Brownian motion model** The geometry model defined in our study can be induced by the following Brownian motion formulation. Taking color as an example, we assume the true color $\theta_0$ is encoded by a continuous function $m(\theta)$, where the function $m(\theta)$ represents the sample path of a Brownian motion with drift, evolving according to the equation

$$dm(\theta) = f(\theta - \theta_0)d\theta + \sigma dB(\theta) \tag{14}$$

from an initial condition $m(0)$, where $f(\theta)$ is a deterministic function, and $B(\theta)$ is a standard Brownian motion which characterizes the Gaussian noise with variance $\sigma^2$. We can think of the increment in the variable $m(\theta)$ over the interval from $\theta_0$ to $\theta_1$ as measuring cumulative activity by the neural channels with preferred colors between $\theta_0$ and $\theta_1$. The von Mises function $f(\theta)$ indicates the shape of the tuning function for each channel. Define $h(\theta - \theta_0) = \int f(\tilde{\theta} - \theta)f(\tilde{\theta} - \theta_0)d\tilde{\theta}$. For any two stimuli, the Euclidean distance between their representation can be expressed as $d(\theta_1, \theta_2)^2 = 2[h(0) - h(\theta_1 - \theta_2)]$. This Brownian motion model induces a geometry of the kind assumed by our model above with the appropriate symmetry. Bayesian estimation on this model and our geometry model is equivalent.

**Population coding model** Our geometry model can also be induced by specifying shift-invariant tuning curves with additive Gaussian noise that are equally spaced on the color wheel, similar to the population coding model with Gaussian noise [33]. The population coding model, further assuming additive Gaussian noise for each neuron with the same magnitude, can be seen as a discrete approximation of the Brownian motion model with finite number of neurons. Importantly, studies based on population coding model argued that Poisson noise was the key factor in determining the shape of the error distributions. In contrast, we propose that the geometry, rather than the Poisson variability, determines the error distributions– a major departure from previous work. In our framework, there are infinitely many ways to specify a particular geometry using tuning curves, as any shift or rotation of manifold will change the tuning curves but lead to an equivalent geometric structure (see Fig. S1) and thus error distribution . The relationship between tuning curves and representational geometry is discussed recently in [48]. Our theory leads to a much compact understanding on how the characteristics of encoding affect error distributions: only changes of tuning curves that modify the shape of the RD function would change the error distributions.

**Psychological scaling model** TCC model proposed in [36] was based on the idea of psychological scaling. In TCC model, on every trial a matching signal is generated for each possible option. The strength of the matching signal is determined by a psychological scaling function, corrupted by Gaussian additive noise. The scaling function is inferred based on an additional set of measurements in triad or quad tasks, in which the subjects perform comparisons on the similarities between two pairs of stimuli. On every trial, the observer's behavioral report is assumed to reflect the option with the largest matching signal. The basic version of TCC model assumes independent noise of the matching signals. In addition, a modified model with correlated noise was also proposed, in which the noise covariance structure was specified independent of the shape of the psychological scaling function. The covariance structure was assumed to reflect the perceptual confusability measured in a separate perceptual task.

Our theory could be connected and compared to the TCC model. First, similar to the TCC model, the likelihood function $L(\theta)$ in our model could be interpreted as a "similarity signal" for each stimulus $\theta$. To this end, the MAP estimate is formally equivalent to selecting a response corresponding to the stimulus with the largest "similarity signal". Our similarity function $g(\theta - \theta_0)$ plays a similar role as the "psychological scaling function" postulated by the TCC model. In both models, this function is independent of the magnitude of the noise. Importantly, our model provides an explanation for the shape of this function through the representational geometry of the encoding, and why it should be unaffected by then noise level. Second, our model predicts that the noise added to the matching signals should follow a Gaussian distribution with a shared variance for each of the candidate responses $\theta$, as posited by the TCC model. Third, our model predicts that the covariance structure of the noisy matching signals should have a cyclical structure, and the covariance function should be a multiple of the similarity function, thus makes a tighter prediction than the TCC model. Critically, the noise correlation in our model exhibits a long-range structure, different from the local structure assumed in the TCC model [60].

[36] showed that TCC model with correlated noise could account for error pattern in 180-AFC and 360-AFC task based 2-AFC data. Importantly, this result relies on assuming a particular noise correlation structure measured in a perceptual matching task. Two remarks are worth mentioning. First, the independent TCC model could not explain these data, due to that the amount of information in the matching signal would increases with the number of options under independent noise. Second, for the correlated TCC model, different noise correlation structure would lead to different error distributions for n-AFC task given the same 2-AFC data. In comparison, the correlation structure in our model is generically constrained by the similarity function and has a more global structure, unlike in TCC model which has a local structure (specified by perceptual confusion matrix in [36]). It is interesting that both model is able to reasonably account for the n-AFC data given the markedly different noise correlation structure– a question requires further study in future.

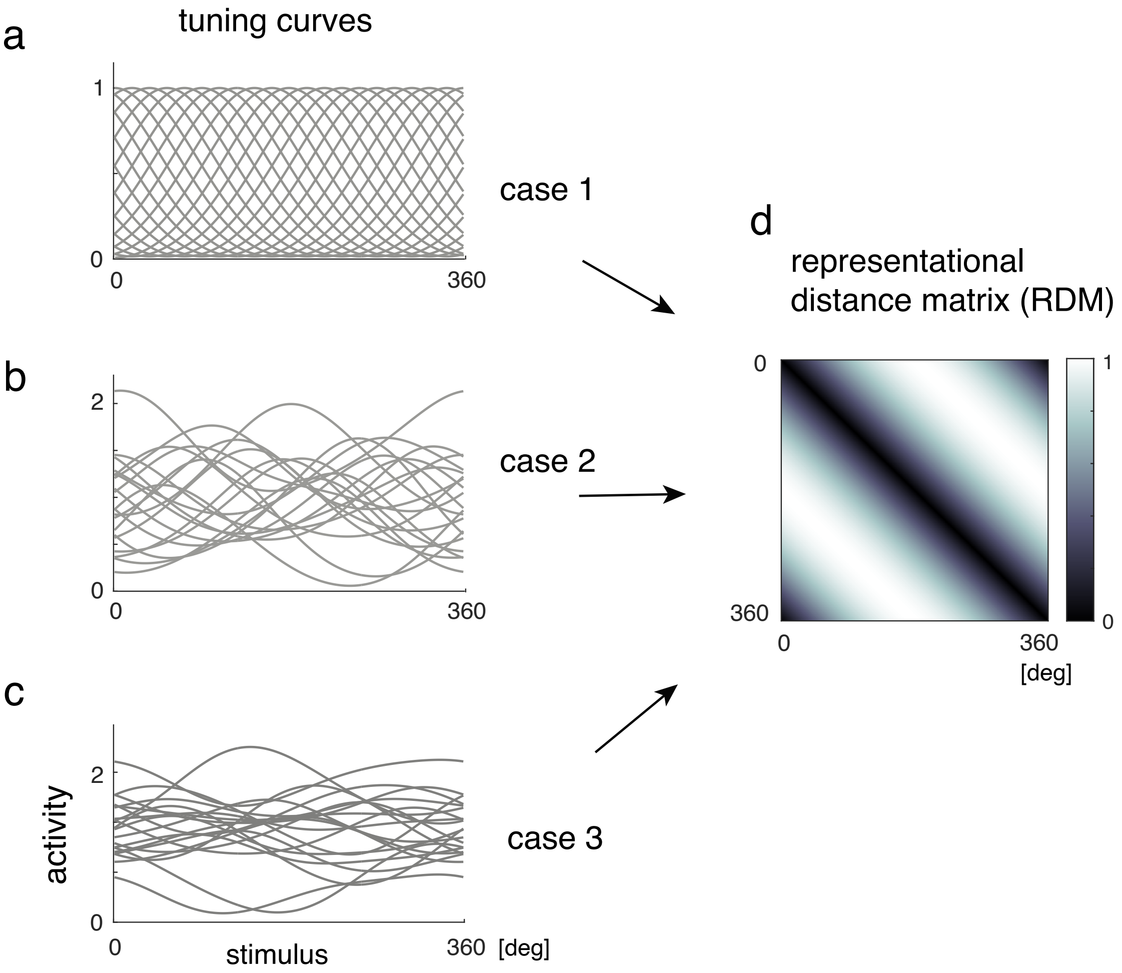


**Figure S1:** Different sets of tuning curves correspond to the same representational geometry. All three tuning configurations correspond to exactly the same geometry. We started from a homogeneous neural population with 20 neurons, and rotate (randomly) and shift the representation to obtain the other two sets of tuning curves.

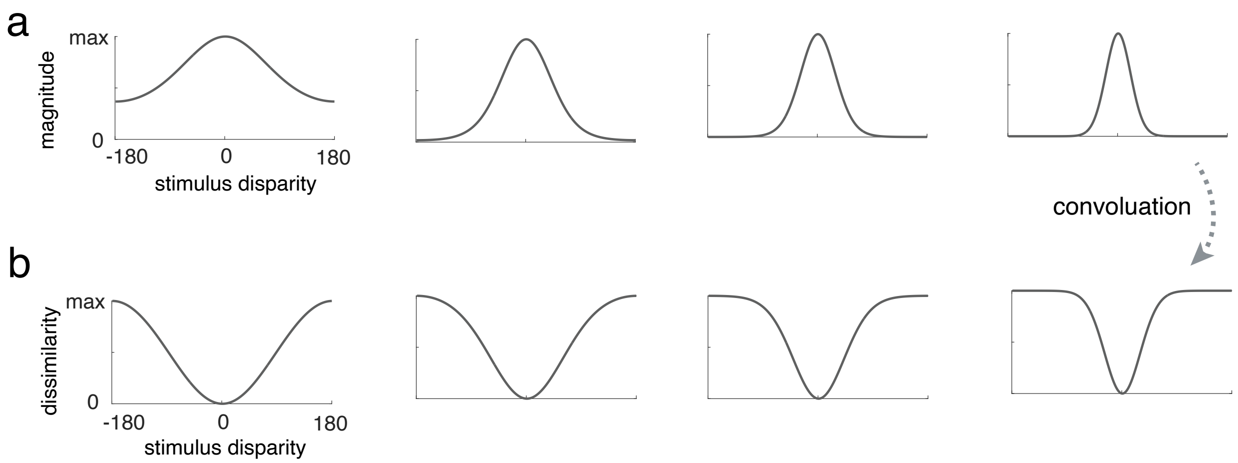


**Figure S2:** Relationship between the kappa ($\kappa$) parameter and representational dissimilarity function. (a) Four different von moises functions, $f()$, with different kappa parameters. (b) The corresponding representational dissimilarity function, which is calculated as $g(\theta - \theta_0) \propto \int f(\tilde{\theta})^2 d\tilde{\theta} - \int f(\tilde{\theta} - \theta) f(\tilde{\theta} - \theta_0) d\tilde{\theta}$. Larger kappa values lead to faster saturation of representational dissimilarity function $g()$.

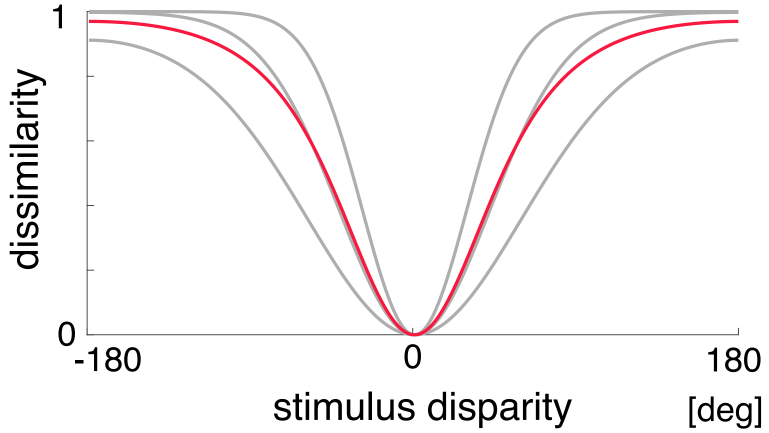


**Figure S3:** Parameterizing the dissimilarity function $g$ using a mixture. In principle, one could obtained different geometry by mixing geometries defined by von Mises functions $f$. Each of the three curves in grey represent a dissimilarity function based on a von Mises function $f$. Red is a mixture of the three by linearly weighting the three grey curves with equal weights. The three grey curves correspond to the dissimilarity function induced by $\kappa = 2, 4, 8$ respectively. Note that, in the paper, we have focused on the geometry induced by von Mises functions. We found even the simple paramterization with one parameter is able to well capture the pattern in the data.

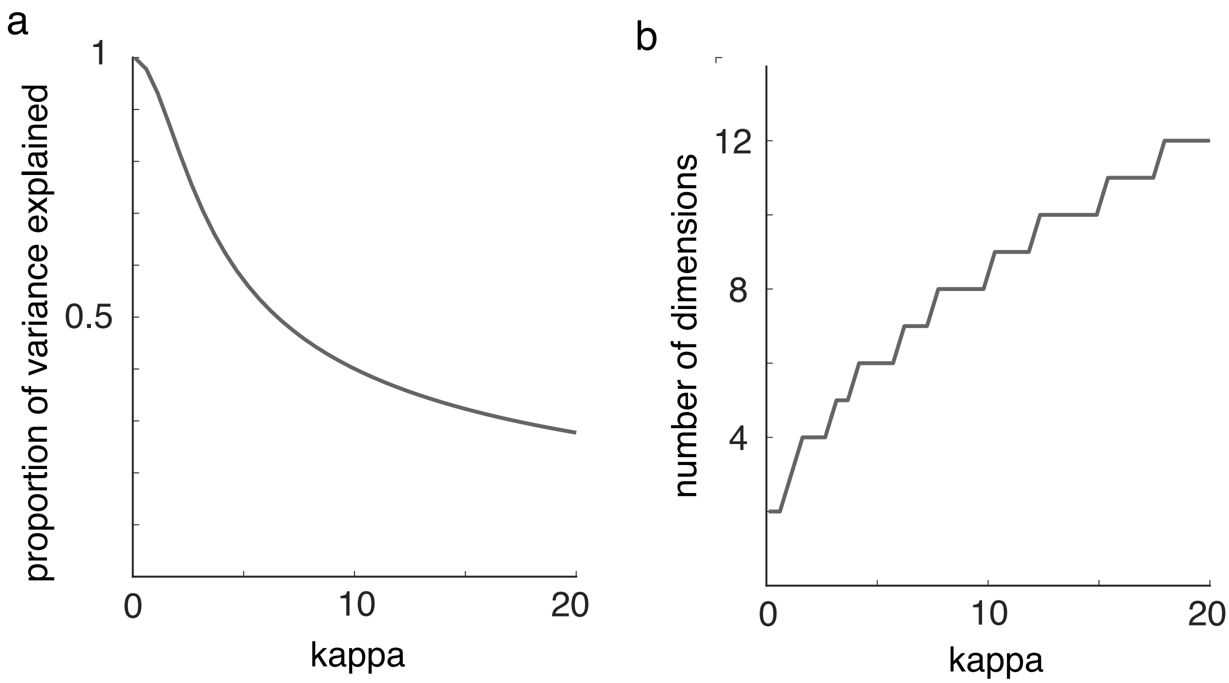


**Figure S4:** Dimensionality. The first panel shows the variance explained by the first two principal dimensions. The second panel shows how many dimensions are needed to explain 95 percent of the variance.

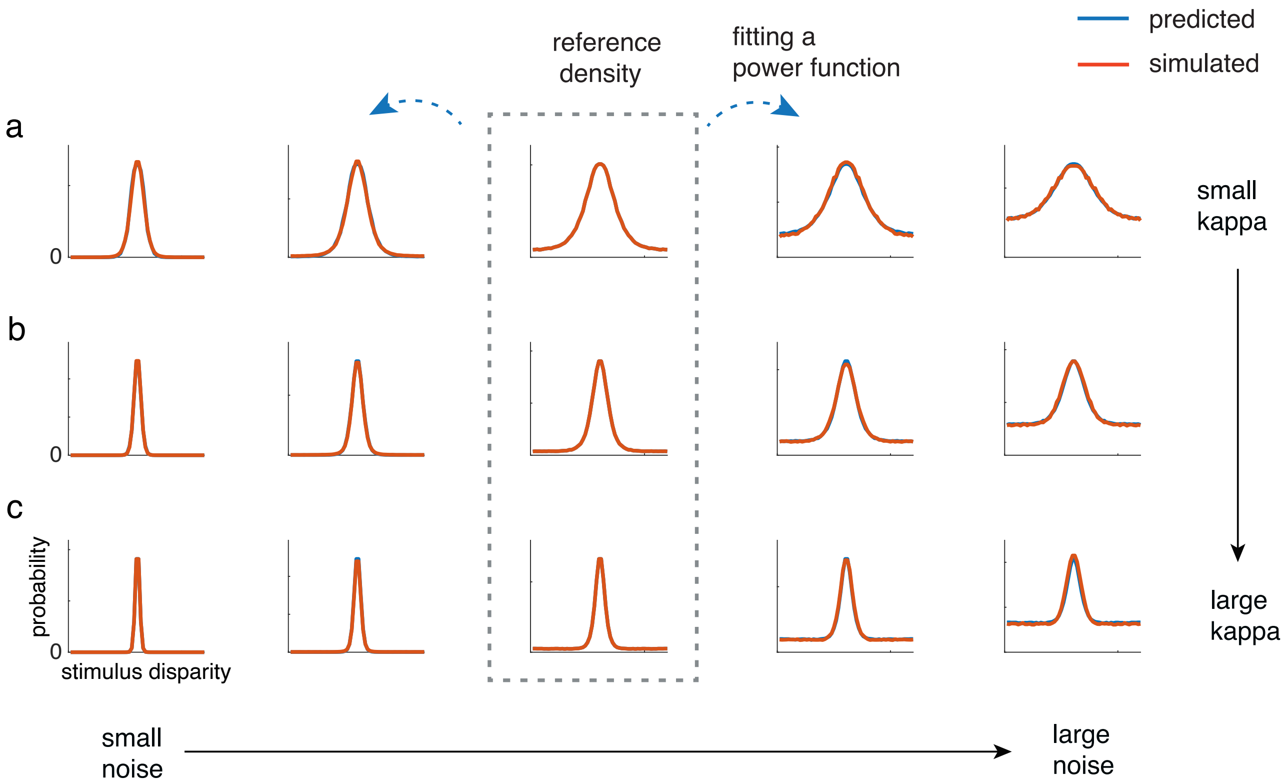


**Figure S5:** The power-law relationship between error densities corresponding to different noise levels when the geometry is fixed.. Each row represents simulations based on a particular kappa ($\kappa$) values ($\kappa = 1, 3, 8$) respectively, each with 5 different noise levels (corresponding to different set sizes). For each row, we use a power function of the reference error distribution (shown in the middle panel) to fit the rest. The results support the power-law relationship between the error distributions corresponding to different noise levels for a fixed $\kappa$.

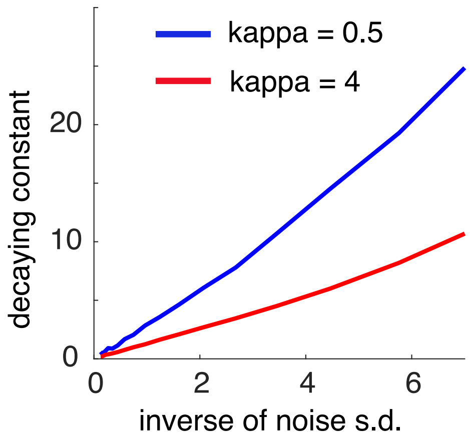


**Figure S6:** The relationship between the inverse of noise s.d. ($\sigma$) and the decaying constant $C$ in Eq. 3. Blue: $\kappa = 0.5$; Red: $\kappa = 4$. The relationship between the two is close to linear when the inverse of the noise is small. The relationship depends on the geometry of the representation.

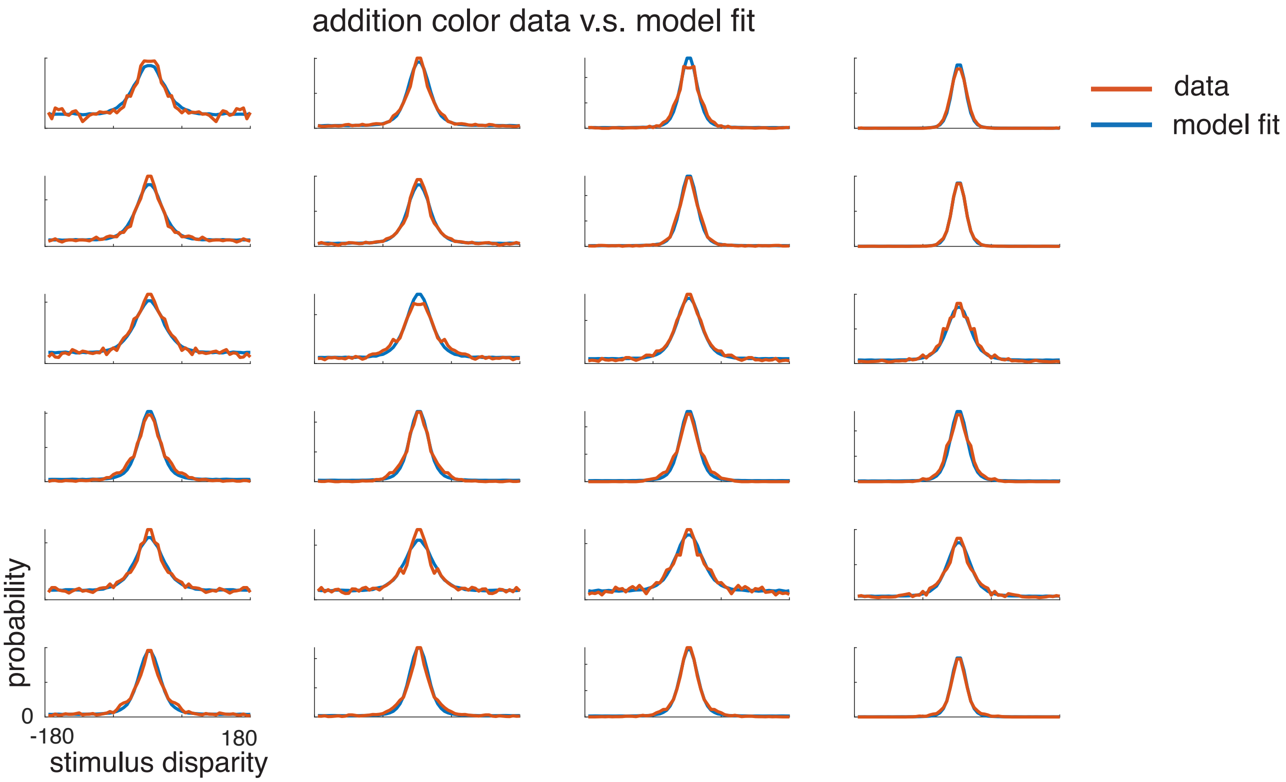


**Figure S7:** Fit of the Bayesian observer model to additional data from color VSTM experiments using continuous report paradigms. Data from [17, 32, 25].

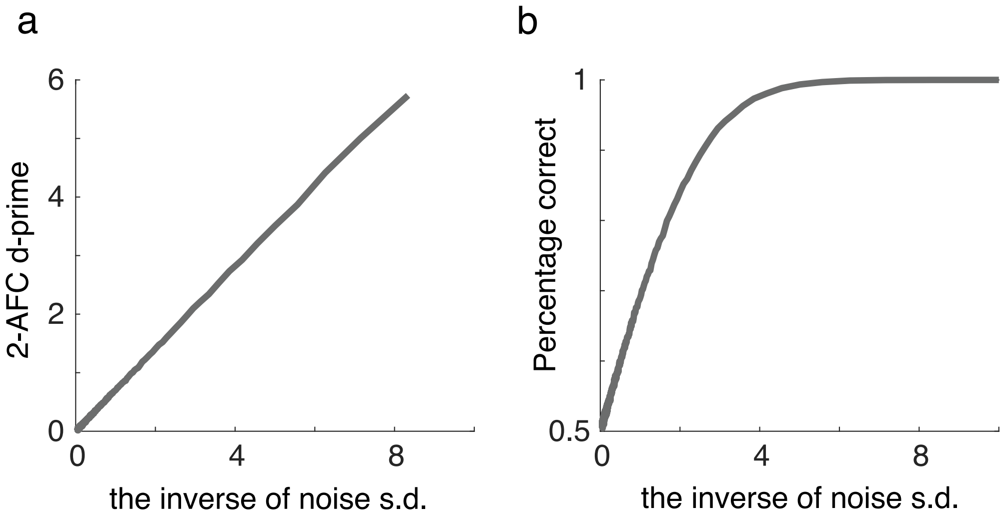


**Figure S8:** (a) The relationship between $1/\sigma$ and d-prime in a 2-AFC task. (b) The relationship between $1/\sigma$ and the percentage correct in a 2-AFC task.
